# Accurate prediction of genetic circuit behavior requires multidimensional characterization of parts

**DOI:** 10.1101/2020.05.30.122077

**Authors:** Galen Dods, Mariana Gómez-Schiavon, Hana El-Samad, Andrew H. Ng

**Author notes:** Correspondence should be addressed to A. H. N. or H. E.-S. These authors contributed equally to this work (Galen Dods, Mariana Gómez-Schiavon).

## Abstract

Mathematical models can aid the design of genetic circuits, but may yield inaccurate results if individual parts are not modeled at the appropriate resolution. To illustrate the importance of this concept, we study transcriptional cascades consisting of two inducible synthetic transcription factors connected in series. Despite the simplicity of this design, we find that accurate prediction of circuit behavior requires mapping the dose responses of each circuit component along the dimensions of both its expression level and its inducer concentration. With such multidimensional characterizations, we were able to computationally explore the behavior of 16 different circuit designs. We experimentally verified a subset of these predictions and found substantial agreement. This method of biological part characterization enables the use of models to identify (un)desired circuit behaviors prior to experimental implementation, thus shortening the design-build-test cycle for more complex circuits.

## Introduction

Synthetic biology utilizes biological parts such as transcription factors to build circuits that perform useful signal processing functions [1,2]. Advancements in DNA synthesis technology have rapidly grown the library of biological parts, but the construction of predictably performing circuits has lagged behind [3]. This lag is due in large part to two factors. First, it is now faster to build new DNA constructs than to characterize them experimentally, leading to the creation of many poorly characterized biological parts [4]. Second, simple phenomenological models of individual parts often fail to predict the behavior of circuits composed of these parts, even in the absence of contextual effects [5] or retroactivity [6]. Building more useful mathematical models of biological parts would greatly facilitate the forward design of genetic circuits with predictable behavior [7–10].

A common feature of genetic circuits is the use of inducible synthetic transcription factors (iSynTFs) [11–13] as facile input nodes that can activate downstream elements in a dose-responsive manner. In *Saccharomyces cerevisiae*, a common architecture for iSynTFs consists of a fusion of a DNA binding domain (DBD), human hormone receptor (HR), and activating domain (AD) [11, 14–16]. Absent their corresponding hormones, these iSynTFs are sequestered in the cytosol via interaction of the HR with Hsp90 [17,18]. This interaction inhibits nuclear localization until hormone is added, enabling dose-responsive control of transcription from a cognate promoter. iSynTFs are an indispensable part of the synthetic biology toolbox. Circuits containing iSynTFs have been used to probe the behavior of synthetic degradation-based feedback [19,20], investigate noise in transcription [11], and study the topology of endogenous circuits [15,21,22].

iSynTFs are commonly characterized via their inducer dose response for one expression level of the transcription factor, but this represents only one dimension of their functionality. Genetic circuits often perform computation by modulating the expression level of transcription factors in a network. Thus, accurate prediction of circuit behavior should be contingent on understanding the behavior of these inducible transcription factors as they change expression level within a circuit.

In this work, we developed a model to predict the behavior of a simple genetic circuit: a transcriptional cascade consisting of two iSynTFs in which the first iSynTF activates expression of a second iSynTF. A simple Hill model reproduced the inducer dose response of an iSynTF at a single expression level, but failed at different iSynTF expression levels due to nonlinearities in the behavior of these biological parts. We overcame this challenge by developing a mechanistic model to account for these nonlinearities and by fitting this model using a two-dimensional inducer and expression level dose response characterization. With this multidimensional characterization, we were able to predict the relationship between inducer concentration and expression level for three different iSynTFs. These models enabled the computational exploration of the full design space of two-step transcriptional cascades, totalling 16 possible circuits. We experimentally validated these simulations for a subset of circuits, confirming the predictive power of the model. These results serve as an example of the type of multidimensional biological part characterization that is required to accurately predict genetic circuit behavior.

## Results and Discussion

To predict the behavior of iSynTFs in genetic circuits, we attempted to fit a simple Hill model to the hormone dose response of an iSynTF in isolation. We first studied GEM, a previously described iSynTF that consists of the Gal4 DBD, estrogen HR, and Msn2 AD, which activates transcription from the pGAL1 promoter in response to estradiol (E2) [11]. We constitutively expressed GEM from pRNR2—a medium strength constitutive promoter previously characterized in the yeast toolkit (YTK) [23]—and measured its dose response, as quantified by the fluorescence output of pGAL1-yellow fluorescent protein (YFP) as a function of E2. A simple Hill model accurately reproduced the basal activity, output saturation, and curvature of this pRNR2-GEM dose response (Fig. 1A; see Methods).

**Figure 1:**
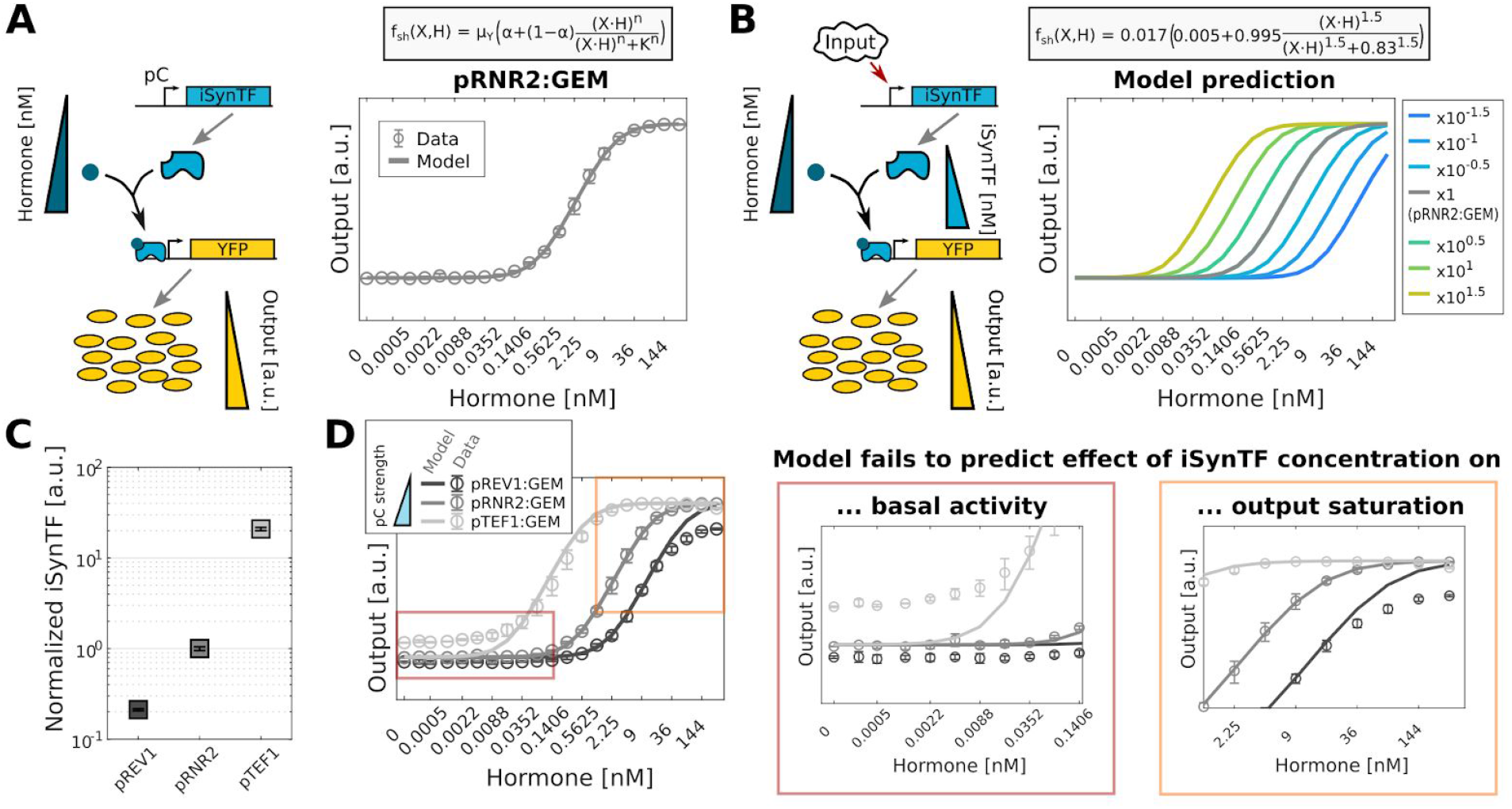
Simple Hill model fit to a single hormone dose response fails to capture the full behavior of iSynTF. **(A)** Left, A constitutively expressed (pC, constitutive promoter) inducible synthetic transcription factor (iSynTF) is bound by its hormone inducer and activates transcription of a downstream YFP reporter (output). Right, Inducer dose response of GEM at a single expression level (pRNR2, constitutive promoter) as a function of hormone, here estradiol. A simple Hill model (see inset: μ_*Y*_, maximum synthesis rate, α, basal activity level; *X*, iSynTF concentration; *H*, hormone concentration; *K*, activation coefficient; *n*, Hill coefficient) was fit to the observed data. **(B)** Left, Expression level of iSynTF can change in response to inputs in a genetic circuit. Right, Simple Hill model prediction (inset for used parameter values) of inducer dose response for different expression levels of GEM (see legend for fold-change values). **(C)** Measurement of constitutive promoter expression levels using a pC-YFP fusion (where pC represents pREV1, pRNR2, or pTEF1). **(D)** Comparison of model predictions and experimental data for GEM inducer dose response at three different expression levels of GEM. Insets in Red and Orange boxes highlight the differences in basal activity and output saturation. Solid lines represent model predictions, open circles and filled squares represent experimental mean, and error bars represent s.d. of three biological replicates.

The output of GEM is dependent on its ligand input, but this hormone dose response relationship may be modulated in non-trivial ways by the expression level of GEM itself. This effect could become significant if GEM is used in a circuit in which its expression level changes. We therefore next sought to understand the relationship between GEM expression level and its hormone dose response. Using the simple Hill model fit to the pRNR2:GEM data, we simulated the dose response of GEM at multiple expression levels around pRNR2 (Fig 1B). Changing the GEM expression level (represented by X in the simple Hill model) simply changed the sensitivity (the half-max point of the sigmoidal curve) of the hormone dose response curve, while maintaining the same basal activity, output saturation, and curvature.

We experimentally tested this prediction by measuring the dose response of GEM at several different expression levels using promoters of different strengths picked from the YTK part library [23]. We selected two promoters, pREV1 and pTEF1, that have lower and higher expression levels than pRNR2, and confirmed their relative expression levels using a promoter fusion to YFP (Fig. 1C). We then used these promoters to drive expression of GEM and experimentally measured each hormone dose response. Contrary to the prediction of the simple Hill model, changing the expression level of GEM did not just shift the hormone dose response sensitivity (Fig. 1D). We also observed a direct effect of GEM expression level on the basal activity (Fig. 1D, red highlight), the output saturation (Fig. 1D, orange highlight), and the curvature of each hormone dose response. These results demonstrate that iSynTFs are dose responsive in two dimensions: hormone concentration and iSynTF expression level. This prompted us to re-examine the choice of model and data used to fit the model.

We hypothesized that the Hill model predictions failed because this simple model did not have sufficient resolution to describe the nonlinear effect of expression level on iSynTF behavior. Furthermore, we hypothesized that we used insufficient data to fit the original model. To address the former, we constructed a mechanistic model that takes into account the allosteric activation of the iSynTF [24], as well as the saturation on the promoter occupancy and elongation rate [25] (Fig. 2A; see Methods). To address the latter, we fit the mechanistic model with the hormone dose responses of GEM at three different expression levels: pREV1:GEM, pRNR2:GEM, and pTEF1:GEM. This new model was able to recapitulate all three of the experimental hormone dose responses (Fig. 2A), and predicted a clear relationship between GEM expression level and the basal activity, output saturation, and curvature (Fig. 2B).

**Figure 2:**
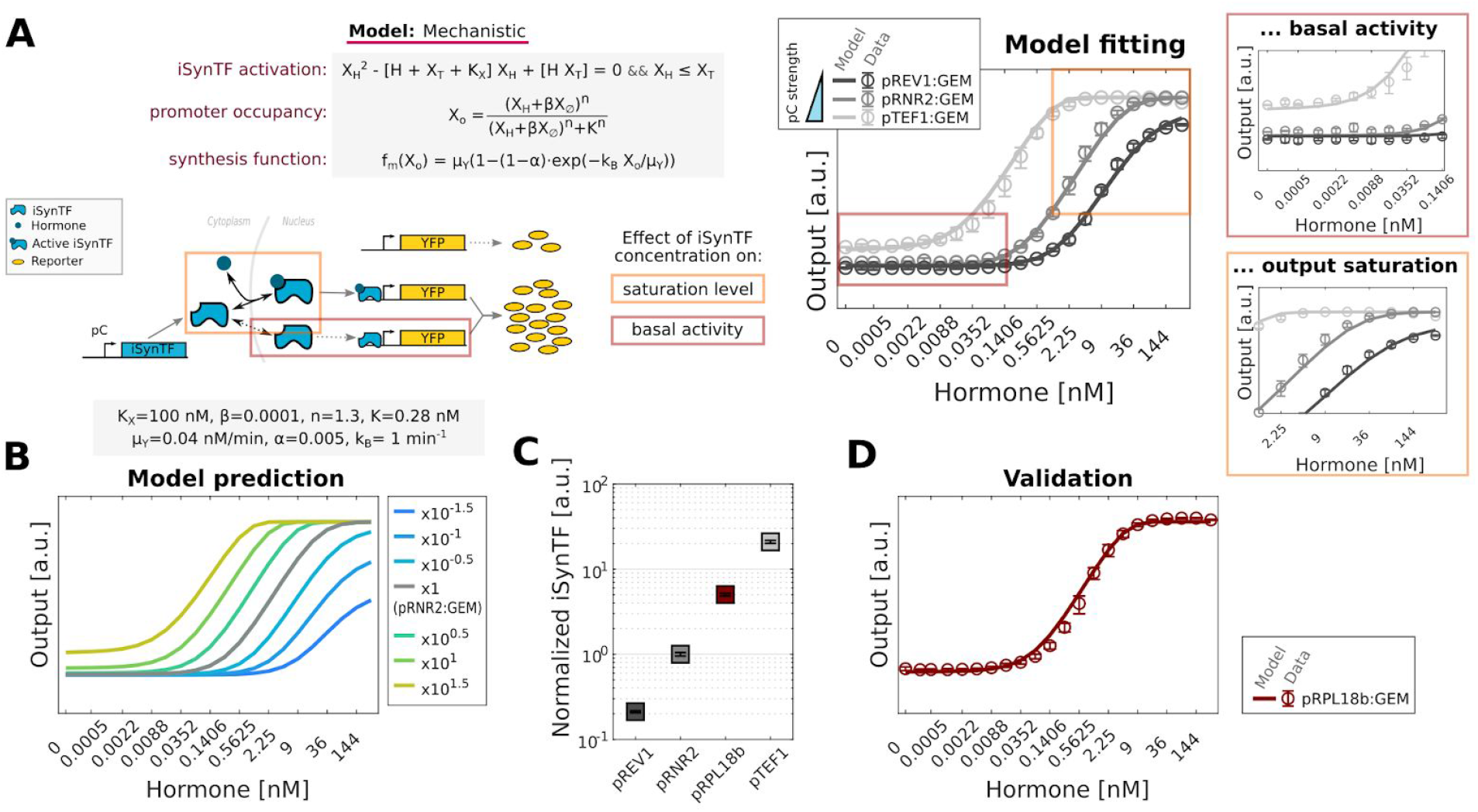
Mechanistic model fit to hormone dose responses at multiple iSynTF expression levels enables accurate prediction of part behavior. **(A)** Left, Schematic of a mechanistic model where iSynTF activation, promoter occupancy, and synthesis dynamics are considered (see Methods for details). Right, Inducer dose response of GEM at three expression levels (pRNR2, pRNR2, and pTEF1 constitutive promoters). The mechanistic model (described in the left) was fit to the observed data. Insets in Red and Orange boxes highlight the recapitulation of the basal activity and output saturation (compare to Fig. 1D). **(B)** Mechanistic model prediction (inset for used parameter values) of inducer dose responses for different expression levels of GEM (see legend for fold-change values). **(C)** Measurement of constitutive promoter expression levels using a pC-YFP fusion including pRPL18B (where pC represents one of pREV1, pRNR2, pRPL18B, or pTEF1). **(D)** Comparison of model prediction and experimental data for pRPL18B:GEM inducer dose response as validation. Solid lines represent model predictions, open circles and filled squares represent experimental mean, and error bars represent s.d. of three biological replicates. See Supplementary Figure 1 for the equivalent analysis using the Z3PM and Z4EM iSynTFs.

To test the accuracy of the mechanistic model, we selected a constitutive promoter of intermediate expression level from the YTK part library, pRPL18B, to drive expression of GEM. We measured the pRPL18B expression level relative to pREV1, pRNR2, and pTEF1 via a YFP promoter fusion (Fig. 2C, red) and input this information into the mechanistic model to predict the dose response of pRPL18B:GEM. Gratifyingly, we found that the model accurately reproduced the basal activity, output saturation, and curvature of the experimental pRPL18B:GEM hormone dose response on which it was not trained (Fig. 2D).

To generalize these results beyond GEM, we examined two other iSynTFs: Z3PM (a fusion of the <u>Z</u>if268 DBD, <u>p</u>rogesterone HR, and <u>M</u>sn2 AD) and Z4EM (a fusion of the <u>Z</u>4 synthetic zinc finger DBD, <u>e</u>strogen HR, and <u>M</u>sn2 AD) [15]. Z3PM activates transcription from pZ3 in a dose responsive fashion to progesterone (Pg), and Z4EM activates transcription from pZ4 in a dose responsive fashion to estradiol (E2). To characterize these iSynTFs, we repeated the workflow developed for GEM: we expressed Z3PM and Z4EM from pREV1, pRNR2, and pTEF1, experimentally measured each hormone dose response, and used these data to fit specific parameters for each iSynTF to the same mechanistic model as above (Fig. S1A). Using the fitted models, we next simulated the effect of iSynTF concentration on the hormone dose response (Fig. S1B). Simulations of Z3PM and Z4EM displayed similar trends to GEM, but they showed a much greater effect of iSynTF expression level on the basal activity and output saturation. Lastly, we validated the accuracy of the Z3PM and Z4EM models against the pRPL18B expression level dose response (Fig. S1C). The Z4EM model accurately captured the basal activity, output saturation, and shape of the pRPL18B dose response curve.The Z3PM model reproduced the output saturation, but it underestimated the basal activity and overestimated the sharpness of the curve.

When comparing the model fittings for each iSynTF, we found that multiple parameter sets fit the observed data equally well, producing similar dose response profiles (Fig. S2). The 100 best fitting parameter sets for each iSynTF showed that some kinetic parameter values were very well constrained (e.g. basal activity α), while others appeared undetermined (e.g. hormone:iSynTF affinity constant, *K_X_*). It may be possible to further resolve differences between the simulations and experiments with a more detailed description of the hormone regulation.

Using these multidimensional, fitted, mechanistic models of GEM, Z3PM, and Z4EM, we explored all possible variants of a two-step transcriptional cascade: a circuit configuration where the constitutively expressed first iSynTF induces expression of a second iSynTF, which in turn induces expression of a YFP reporter (Fig. 3A). With two orthogonal HRs, there are four possible configurations of the three iSynTFs (GEM → Z3PM, Z4EM → Z3PM, Z3PM → GEM, Z3PM → Z4EM). Taking into account four possible expression levels for the first iSynTF (pREV1, pRNR2, pRPL18B, pTEF1), in total there are sixteen possible circuit variants. Because GEM, Z3PM, and Z4EM each have a unique response to hormone and changing expression level, we expected that each circuit variant would behave differently in response to the two hormone inducers. In agreement, the simulations displayed different responses to both inducers, basal activities, and output saturations (Fig. S3). These multidimensional, fitted models enable efficient screening of these circuit variants, guiding the selection of designs to be tested experimentally.

**Figure 3:**
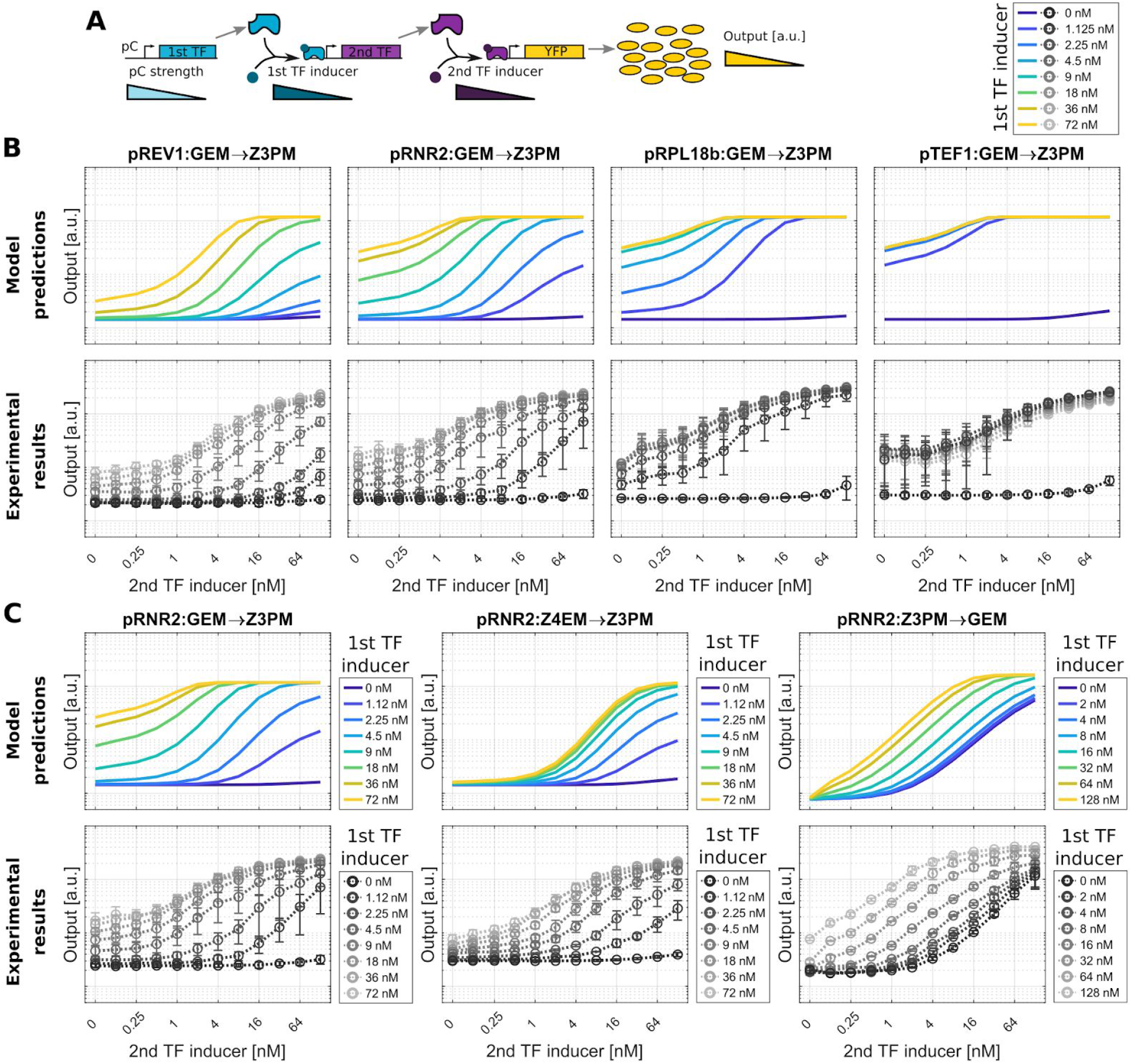
Using refined models to explore circuit designs. **(A)** A constitutively expressed (pC, constitutive promoter) iSynTF (1st TF) is bound by its hormone inducer (1st TF inducer) and activates transcription of a second downstream iSynTF (2nd TF), which in turn binds its hormone inducer (2nd TF inducer) and activates transcription of the downstream YFP reporter (output). **(B)** Comparison of the mechanistic model predictions (top row) and experimental data (bottom row) for circuit output as a function of 2nd TF inducer (x-axis, Pg) at four different expression levels of the 1st TF (see plot titles). The 2nd TF inducer dose responses were simulated or measured at eight different concentrations of the 1st TF inducer (see legend in the top). **(C)** Analogous to panel B, but varying the circuit configuration instead of the promoter strength (see plot titles). Solid lines represent model predictions, open circles and dotted lines represent experimental mean, and error bars represent s.d. of three biological replicates. See Supplementary Figure 3 for simulations of all circuit designs.

We sought to verify the accuracy of the model simulations by experimentally measuring the hormone dose responses of a subset of circuit variants. First, we studied the effect of changing the first iSynTF expression level for a single configuration by measuring the output of GEM → Z3PM at all four expression levels of GEM (Fig. 3B). We found that the model accurately predicted several key aspects of the circuit behavior such as the changing output saturation and curvature. Notably, the simulations underestimated the effect of E2 on the basal activity in the absence of Pg. For comparison, we simulated the output of these circuits using the simple Hill model fit only to the pRNR2:iSynTF dose responses (Fig. S4A) or the mechanistic model fit only to the pRNR2:iSynTF dose responses (Fig. S4B). The predictions with the simple Hill model show no change in output saturation or basal activity (Fig. S4D), while the predictions with the mechanistic model with no multidimensional characterization show excessive basal activity and leaky expression (Fig. S4E). Neither match the experimental results, indicating that the multidimensional characterization is necessary for the accurate prediction of circuit behavior.

Next, we compared three circuit configurations (GEM → Z3PM, Z4EM → Z3PM, Z3PM → GEM) at the pRNR2 expression level of the first iSynTF (Fig. 3C). As predicted by the model simulations, the Z3PM → GEM configuration displayed the greatest responsiveness to the second TF inducer in the absence of the first TF inducer. The simulations were also able to qualitatively predict the curvature of the second TF inducer dose response curves, as well as the effect of the first TF inducer on output saturation. As before, the simulations underestimated the effect of the first TF inducer on circuit output in the absence of the second TF inducer.

This slight quantitative discrepancy can likely be explained by a shortcoming in the model’s ability to predict the expression level of the second TF as a function of the first TF inducer. The model assumes that expression of the second TF will be equivalent to expression of a YFP reporter, despite the fact that contextual factors such as transcript length (e.g. YFP vs. iSynTF), 5’ UTR, or terminator sequence (e.g. tPGK1 vs. tSSA1) have a known effect on output [26–28]. Unfortunately, directly measuring the expression level of the second iSynTF via a fluorescent protein fusion is challenging, as this also alters these variables. Despite this shortcoming, the simulations were still able to predict key qualitative aspects of the experimental data based on the circuit configuration and expression level. Taken together, these data indicate that models can serve as a guide to genetic circuit design when an appropriate characterization of individual parts is performed.

Quantitative model fits can be important in certain scenarios, such as building models to automate genetic circuit design. Recently an algorithm was developed to automate the design of genetic logic gates given a set of user constraints and a library of transcriptional repressors [29]. The algorithm was successful at designing most circuits, but was not perfect; failed circuits adopted intermediate states that did not meet the digital threshold as a result of unexpected part behavior. This issue of unpredictable part behavior plagues synthetic biology in general and is a thorn in the side of many modeling efforts.

An alternative application of part models, given the universal issue of unpredictable part behavior, is theory-guided exploration of potential circuit behavior. Theory can reveal circuit topologies that produce a desired phenotype, and has been used in the past to study biochemical adaptation [30,31]. However, insights gained from these studies can often be difficult to translate into actual designs because there is no guarantee that biological parts exist in the required parameter regimes to implement such circuit designs. It may be possible to use our part characterization methodology to constrain the parameter space of theoretical explorations, biasing the results towards circuits that can be constructed using existing parts. However, our results suggest that parts would need to be characterized based on the design goal of the circuit. For example, dynamic part data would need to be collected if dynamic circuit behavior is desired, and functionality of parts under stressors such as glucose depletion may be important if the circuit is expected to function under stress-inducing conditions.

Model based simulation of genetic circuit behavior can guide circuit designs and limit the number of constructs that need to be tested to achieve a desired behavior. In this work, we focused on a two-step transcriptional cascade of iSynTFs where the expression level of the second iSynTF in the cascade changes in response to input from the first iSynTF. We presented a methodology for model design and fitting that considered the effect of both inducer concentration and expression level on iSynTF output, thus enabling the accurate prediction of genetic circuit behavior. Our results highlight the necessity of understanding biological part behavior in the functional context of potential circuit designs. Such multidimensional characterization requires an upfront investment of time, but it can pay dividends in the long-term by shortening the design-build-test cycle for more complex circuits.

## Methods

### Construction of DNA Constructs

Hierarchical golden gate assembly was used to assemble plasmids for yeast strain construction [23]. Individual parts were ordered as gBlocks (IDT) or PCR amplified (NEB Q5 High-Fidelity 2x Master Mix). PCR products were purified with a GeneJET PCR Purification Kit (Thermo Fisher Scientific). These sequences were domesticated with FastDigest Esp3I (Thermo Fisher Scientific). Transcriptional cassettes were constructed using BsaI-HF v2 (NEB). Multigene plasmids were constructed using FastDigest Esp3I. Plasmids are listed in Supplementary Table 1, and oligos are listed in Supplementary Table 2. The pZ4 sequence was modified from McIssac *et al. [15]* to remove a Gal4 binding site (sequence in supplement).

Chloramphenicol and Ampicillin resistant plasmids were transformed into chemically competent Mach1 *E. coli* (QB3 Macrolab), while Kanamycin resistant plasmids were transformed into chemically competent XL1 Blue *E. coli* (QB3 Macrolab). Cultures were grown over the course of the day (Mach1) or overnight (XL1) before prep. Following growth, cultures were prepared using a GeneJET Plasmid Miniprep Kit (Thermo Fisher Scientific). Part plasmids were verified by sequencing (Elim Biopharmaceuticals) using the listed sequencing primers, while all other plasmids were verified by restriction enzyme digestion.

### Yeast Growth Media

Overnight yeast cultures were grown in YPD (1% w/v bacto-yeast extract; 2% w/v bacto-peptone; and 2% w/v dextrose). Yeast transformation cultures were diluted into fresh YPD. Cultures for flow cytometry were diluted into SDC (0.67% w/v Difco yeast nitrogen base without amino acids; 0.2% complete supplement mixture (MP Biomedicals); and 2% w/v dextrose). For prototrophic selection following yeast transformation, SDC agar plates with the appropriate selection were used (Teknova).

### Construction of Yeast Strains

All DNA constructs were transformed into a yeast strain derived from BY4741 *(MATa his3Δ1 leu2Δ0 met15Δ0 ura3Δ0*) that had the HIS3 locus repaired. Yeast transformations were performed as described previously [23] with modifications. One wash with 100 nM lithium acetate was performed. DNA was combined with 115 microliters of transformation mixture and incubated at 42 °C for 30 minutes. All DNA constructs were genomically integrated. Three microliters of prepared plasmid were linearized for integration in a twenty microliter NotI-HF (NEB) reaction for one hour and then added to the transformation mixture without purification. Strains are listed in Supplementary Table 3.

### Flow Cytometry experiments

Yeast strains were streaked out onto YPD plates from glycerol stocks. Individual colonies were picked into 1 mL of YPD in a 2-mL V-bottom 96-well block (Corning/Costar) for overnight growth at 30 °C and 900 rpm in a Multitron shaker (Infors HT). For the individual iSynTFs experiments, overnight cultures were diluted 1:500 in 12 mL of fresh SDC in an 8-row block and 450 microliters were aliquoted into a row across 2 new 96 well blocks. For the cascade experiments, overnight cultures were diluted 1:500 in 45 mL of fresh SDC in a 50 mL trough (Corning) and 400 microliters were aliquoted into all wells of a new 96 well block. The YFP-promoter fusion strains were diluted 1:500 in 500 μL of fresh SDC in a new 96 well block. Following dilution, blocks were returned to the shaker for a 2 hour outgrowth.

During the 2 hour outgrowth, estradiol (Sigma-Aldrich) and progesterone (Fisher Scientific) induction gradients were prepared. Ten-times concentrated solutions were prepared in fresh SDC from 36 micromolar (estradiol) and 32 micromolar (progesterone) stock solutions. Gradients were then prepared by a one-to-one serial dilution from the maximum induction solution. For the individual iSynTF experiments, 50 microliters of the corresponding solution were added to the appropriate wells. For the cascade characterizations, 50 microliters of each solution were added to each well in the corresponding combinations. Blocks were then returned to the shaker for 4 hours.

Following the 4 hour induction, the cultures were prepared for flow cytometry. One hundred microliters of culture were mixed with 100 microliters of fresh SDC in a 96-well U-bottom microplate (greiner bio-one). Samples were measured on a BD LSRFortessa X20 (BD Biosciences) using a high-throughput sampler. YFP-Venus fluorescence was measured using the FITC-H channel (voltage = 434). Measurements were normalized by dividing by SSC-H (voltage = 200). Analysis was performed with Python 3.7, custom scripts, and the FlowCytometryTools package. All experiments were performed in triplicate, with replicates collected on separate days. Reported values represent the mean and standard deviation of median normalized fluorescence values of the triplicates.

### Model: Simple Hill Function

Under this model, we assume that the iSynTF is constitutively produced and has reached its steady state concentration (*X*). The concentration of hormone in the media is denoted by *H*. Then the steady-state concentration of the reporter protein is described as a simple Hill function with maximum synthesis rate μ_*Y*_, basal activity α ∈ [0, 1], dissociation constant *K*, and Hill coefficient *n*:

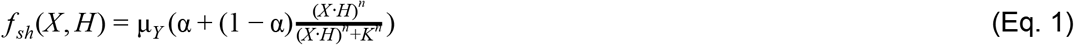

### Model: Mechanistic

As shown in Fig. 1, the simple Hill model described above fails to capture the effect of the iSynTF concentration (*X*) on the basal expression level and saturation of the inducer dose response output. To better recapitulate the observed behavior, we propose a mechanistic model including the following considerations:

1. In the absence of hormone (*H* = 0), increasing iSynTF expression (e.g. using a stronger constitutive promoter) increases the output expression level, suggesting some leaky or basal activation of the regulated promoter by free iSynTF.
2. As the iSynTF expression decreases, a minimum output expression level is observed for low hormone concentrations, suggesting some leaky expression of the regulated promoter independent of the iSynTF.
3. As both iSynTF and hormone concentration increase, the output expression level saturates at a maximum value. This can be explained both by the saturation of the regulated promoter occupancy, and by the saturation of the number of polymerases simultaneously transcribing the output gene.
4. With low iSynTF expression, the output expression level saturates at a lower level as hormone concentration increases, suggesting that the stoichiometric relationship of the iSynTF and hormone molecules might play an important role in output regulation.

Then, the proposed model is:

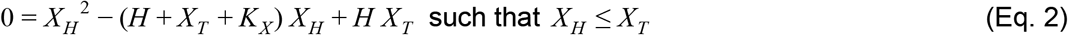

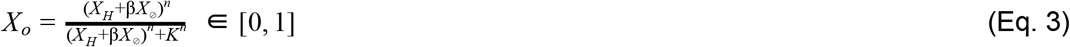

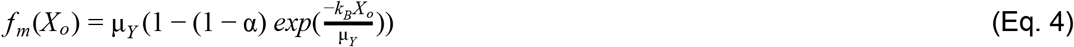

Where *X*_⊘_ is the concentration of free (inactive) iSynTF, *X_H_* is the active iSynTF (i.e. *X*_⊘_ bound to the hormone), and *X_T_* = *X*_⊘_ + *X_H_* is the total iSynTF concentration in the cell (e.g. determined by the used promoter driving iSynTF expression). *H* is the total intracellular hormone concentration, which is assumed constant throughout the experiment, and proportional to the amount added to the media. *X_o_* is the regulated promoter occupancy, which is modeled as a Hill function of the active iSynTF and a fraction (β) of the inactive iSynTF in the nucleus, with Hill coefficient *n* and dissociation constant *K*. Finally, the synthesis rate of the regulated promoter *f_m_*(*X_o_*) is modeled as proposed in Ben-Tabou de-Leon & Davidson (2009), where μ_*Y*_ is the maximum synthesis rate given the translocation rate and gene, *k_B_* is the efficiency rate of the transcription factor, and α ∈ [0,1] is the basal expression of the output gene (in the absence of iSynTF). Here, (1) the parameter β represents the basal activation by free iSynTF; (2) the parameter α represents the leakiness of the regulated promoter; (3) both Eqs. 3 & 4 consider the two possible sources of saturation; and (4) Eq. 2 incorporates the stoichiometric relationship between the free iSynTF, hormone, and active iSynTF. This model recapitulates most of the qualitative behavior of the iSynTF regulation for several constitutive promoter strengths and hormone concentrations (see Fig. 2 & Fig. S1).

### Model: Fitting

The goal is to minimize the error between the observed data (*D*) and the model prediction (*Y*) for a given model and parameter set (*θ*). We define our error function simply as the sum of squared errors in logarithmic scale:

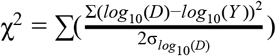

Then we use a Metropolis Random Walk algorithm to explore the parameter space implemented as follows:

1. Choose some initial parameters θ_1_ and calculate its fitting error χ^2^_(1)_.
2. Iterate over *t* = {1, 2,…, *t_MAX_*} as follows:

a. Draw a random proposal ϕ ~ θ_(*t*)_ × 2^*N*_║θ║_(0,Σ)^ where *N*_║θ║_(0, Σ) is a Multivariate Normal distribution with the same dimension as θ_(*t*)_, mean zero and covariance matrix Σ = 0.1. We enforce that the parameters stay in a realistic range with the following limits: *K_X_* = [1 × 10^−4^, 100]; β = [2 × 10^−7^, 0.2]; *n* = [1 × 10^−5^, 10]; *K* = [1 × 10 ^4^, 100]; α = [2 × 10 ^7^,0.2]. And γ_*Y*_ = 0.01*min*^−1^ and μ_*Y*_ = *max*(*D*)· γ_*Y*_ have fixed values.
b. We construct a likelihood function using a Gaussian function:

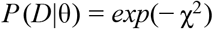

where θ is the set of parameter to be optimized, *D* is the optimal data, and χ^2^ is the error function. Note the likelihood is maximal when the error is minimal. Then we calculate the likelihood ratio:

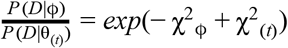 Accept the proposed ϕ if the ratio is larger than a random number ~ *U*[0,1].The proposed value is always accepted if the error is smaller (i.e. it is better).
c. Update parameters θ_(*t*+1)_ ← ϕ with probability 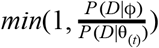; otherwise, θ_(*t*+1)_ ← θ_(*t*)_.

## Author Contribution

G. D., M. G.-S., H. E.-S., and A. H. N. designed the study and experiments. G. D. and A. H. N. performed experiments. M. G.-S., H. E.-S., and A. H. N. developed the computational models. M. G.-S. performed the computational simulations. G. D., M. G.-S., and A. H. N. wrote the manuscript. G. D., M. G.-S., H. E.-S., and A. H. N. edited and approved the manuscript.

## Abbreviations

iSynTF: inducible synthetic transcription factor
YFP: yellow fluorescent protein
GEM: Gal4 DNA binding domain, estradiol ligand binding domain, Msn2 activating domain
Z3PM: Z3 DNA binding domain, progesterone ligand binding domain, Msn2 activating domain
Z4EM: Z4 DNA binding domain, estradiol ligand binding domain, Msn2 activating domain

## Acknowledgements

The authors would like to thank the members of the El-Samad lab for useful comments and discussion on the manuscript. This work was supported by the Defense Advanced Research Projects Agency, Contract No. HR0011-16-2-0045 to H.E.-S and National Science Foundation grant DBI-1548297 to H.E.-S. The content and information does not necessarily reflect the position or the policy of the government, and no official endorsement should be inferred. H.E.-S. is a Chan-Zuckerberg investigator.

## Supporting Information

Supplemental figures; plasmids, oligos, and strains used; pZ4 (-Gal4 site) sequence

## References

1. Elowitz MB, Leibler S. A synthetic oscillatory network of transcriptional regulators. Nature. 2000. pp. 335–338. doi:10.1038/35002125

2. Gardner TS, Cantor CR, Collins JJ. Construction of a genetic toggle switch in Escherichia coli. Nature. 2000;403: 339–342.

3. Purnick PEM, Weiss R. The second wave of synthetic biology: from modules to systems. Nat Rev Mol Cell Biol. 2009;10: 410–422.

4. Kwok R. Five hard truths for synthetic biology. Nature. 2010;463: 288–290.

5. Cardinale S, Arkin AP. Contextualizing context for synthetic biology--identifying causes of failure of synthetic biological systems. Biotechnol J. 2012;7: 856–866.

6. Del Vecchio D, Ninfa AJ, Sontag ED. Modular cell biology: retroactivity and insulation. Mol Syst Biol. 2008;4: 161.

7. Murphy KF, Balázsi G, Collins JJ. Combinatorial promoter design for engineering noisy gene expression. Proc Natl Acad Sci U S A. 2007;104: 12726–12731.

8. Rosenfeld N, Young JW, Alon U, Swain PS, Elowitz MB. Accurate prediction of gene feedback circuit behavior from component properties. Mol Syst Biol. 2007;3: 143.

9. Stricker J, Cookson S, Bennett MR, Mather WH, Tsimring LS, Hasty J. A fast, robust and tunable synthetic gene oscillator. Nature. 2008;456: 516–519.

10. Davidsohn N, Beal J, Kiani S, Adler A, Yaman F, Li Y, et al. Accurate predictions of genetic circuit behavior from part characterization and modular composition. ACS Synth Biol. 2015;4: 673–681.

11. Aranda-Díaz A, Mace K, Zuleta I, Harrigan P, El-Samad H. Robust Synthetic Circuits for Two-Dimensional Control of Gene Expression in Yeast. ACS Synth Biol. 2017;6: 545–554.

12. Mercer AC, Gaj T, Sirk SJ, Lamb BM, Barbas CF 3rd. Regulation of endogenous human gene expression by ligand-inducible TALE transcription factors. ACS Synth Biol. 2014;3: 723–730.

13. Donahue PS, Draut JW, Muldoon JJ, Edelstein HI, Bagheri N, Leonard JN. The COMET toolkit for composing customizable genetic programs in mammalian cells. Nat Commun. 2020;11: 779.

14. McIsaac RS, Silverman SJ, McClean MN, Gibney PA, Macinskas J, Hickman MJ, et al. Fast-acting and nearly gratuitous induction of gene expression and protein depletion in Saccharomyces cerevisiae. Mol Biol Cell. 2011;22: 4447–4459.

15. McIsaac RS, Oakes BL, Wang X, Dummit KA, Botstein D, Noyes MB. Synthetic gene expression perturbation systems with rapid, tunable, single-gene specificity in yeast. Nucleic Acids Res. 2013;41: e57–e57.

16. McIsaac RS, Gibney PA, Chandran SS, Benjamin KR, Botstein D. Synthetic biology tools for programming gene expression without nutritional perturbations in Saccharomyces cerevisiae. Nucleic Acids Res. 2014;42: e48.

17. Pratt WB. The hsp90-based chaperone system: involvement in signal transduction from a variety of hormone and growth factor receptors. Proc Soc Exp Biol Med. 1998;217: 420–434.

18. Smith DF, Toft DO. Minireview: The Intersection of Steroid Receptors with Molecular Chaperones: Observations and Questions. Molecular Endocrinology. 2008. pp. 2229–2240. doi:10.1210/me.2008-0089

19. Ng AH, Nguyen TH, Gómez-Schiavon M, Dods G, Langan RA, Boyken SE, et al. Modular and tunable biological feedback control using a de novo protein switch. Nature. 2019;572: 265–269.

20. Langan RA, Boyken SE, Ng AH, Samson JA, Dods G, Westbrook AM, et al. De novo design of bioactive protein switches. Nature. 2019;572: 205–210.

21. Hackett SR, Baltz EA, Coram M, Wranik BJ, Kim G, Baker A, et al. Learning causal networks using inducible transcription factors and transcriptome-wide time series. Mol Syst Biol. 2020;16: e9174.

22. McIsaac RS, Petti AA, Bussemaker HJ, Botstein D. Perturbation-based analysis and modeling of combinatorial regulation in the yeast sulfur assimilation pathway. Mol Biol Cell. 2012;23: 2993–3007.

23. Lee ME, DeLoache WC, Cervantes B, Dueber JE. A Highly Characterized Yeast Toolkit for Modular, Multipart Assembly. ACS Synth Biol. 2015;4: 975–986.

24. Phillips R, Belliveau NM, Chure G, Garcia HG, Razo-Mejia M, Scholes C. Figure 1 Theory Meets Figure 2 Experiments in the Study of Gene Expression. Annual Review of Biophysics. 2019. pp. 121–163. doi:10.1146/annurev-biophys-052118-115525

25. Ben-Tabou de-Leon S, Davidson EH. Modeling the dynamics of transcriptional gene regulatory networks for animal development. Dev Biol. 2009;325: 317–328.

26. Chen Y-J, Liu P, Nielsen AAK, Brophy JAN, Clancy K, Peterson T, et al. Characterization of 582 natural and synthetic terminators and quantification of their design constraints. Nat Methods. 2013;10: 659–664.

27. Kosuri S, Goodman DB, Cambray G, Mutalik VK, Gao Y, Arkin AP, et al. Composability of regulatory sequences controlling transcription and translation in Escherichia coli. Proc Natl Acad Sci U S A. 2013;110: 14024–14029.

28. Mutalik VK, Guimaraes JC, Cambray G, Lam C, Christoffersen MJ, Mai Q-A, et al. Precise and reliable gene expression via standard transcription and translation initiation elements. Nat Methods. 2013;10: 354–360.

29. Nielsen AAK, Der BS, Shin J, Vaidyanathan P, Paralanov V, Strychalski EA, et al. Genetic circuit design automation. Science. 2016;352: aac7341.

30. Ma W, Trusina A, El-Samad H, Lim WA, Tang C. Defining network topologies that can achieve biochemical adaptation. Cell. 2009;138: 760–773.

31. Briat C, Gupta A, Khammash M. Antithetic Integral Feedback Ensures Robust Perfect Adaptation in Noisy Biomolecular Networks. Cell Syst. 2016;2: 15–26.

