## Supplementary Information for "Accurate prediction of genetic circuit behavior requires multidimensional characterization of parts"

### Supplemental Figures

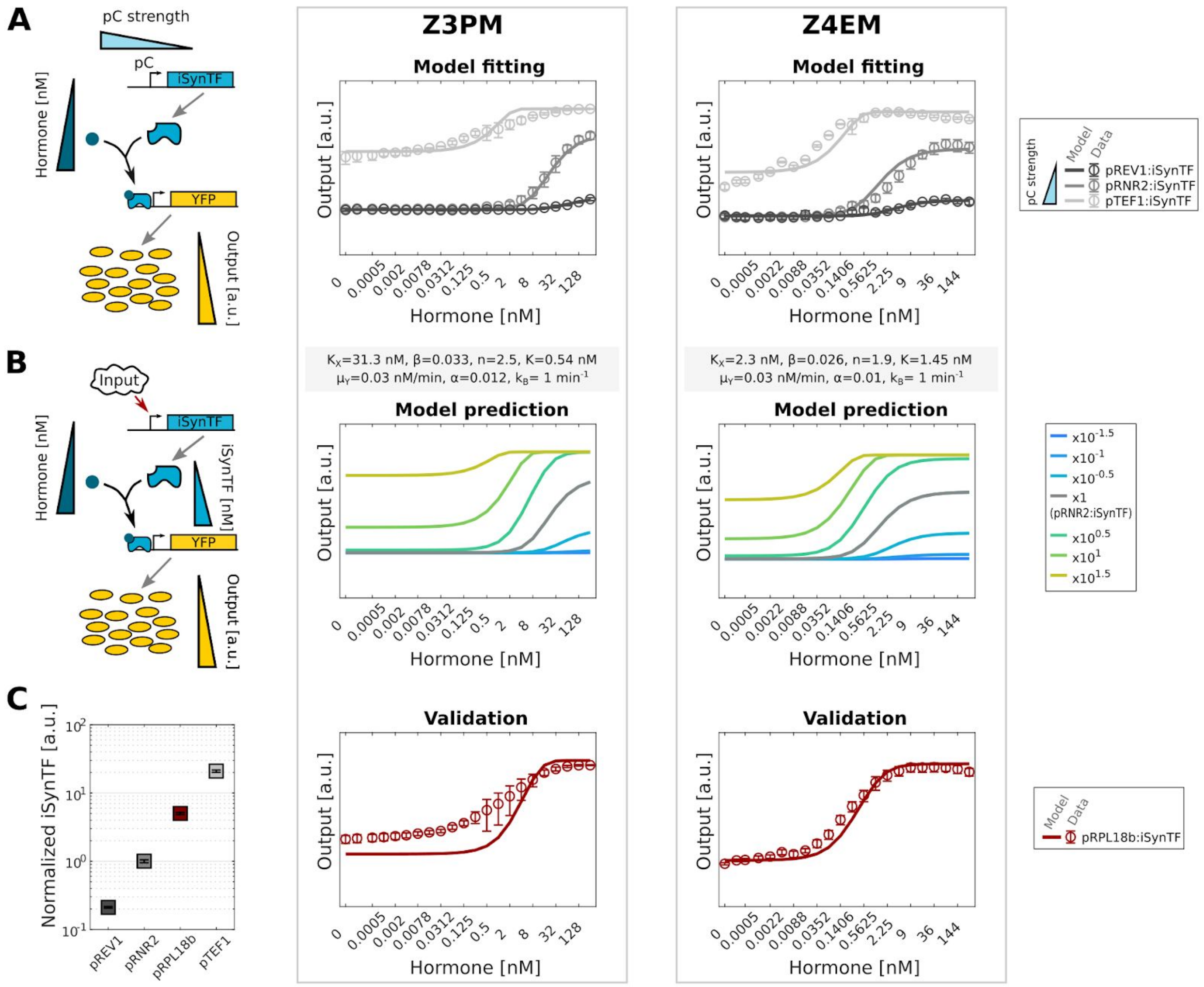


**Supplementary Figure 1: Multidimensional characterization and mechanistic model fit for the Z3PM and Z4EM iSynTFs.** Same figure description as Figure 2. See Methods and Supplementary Figure 2 for parameter fitting details.

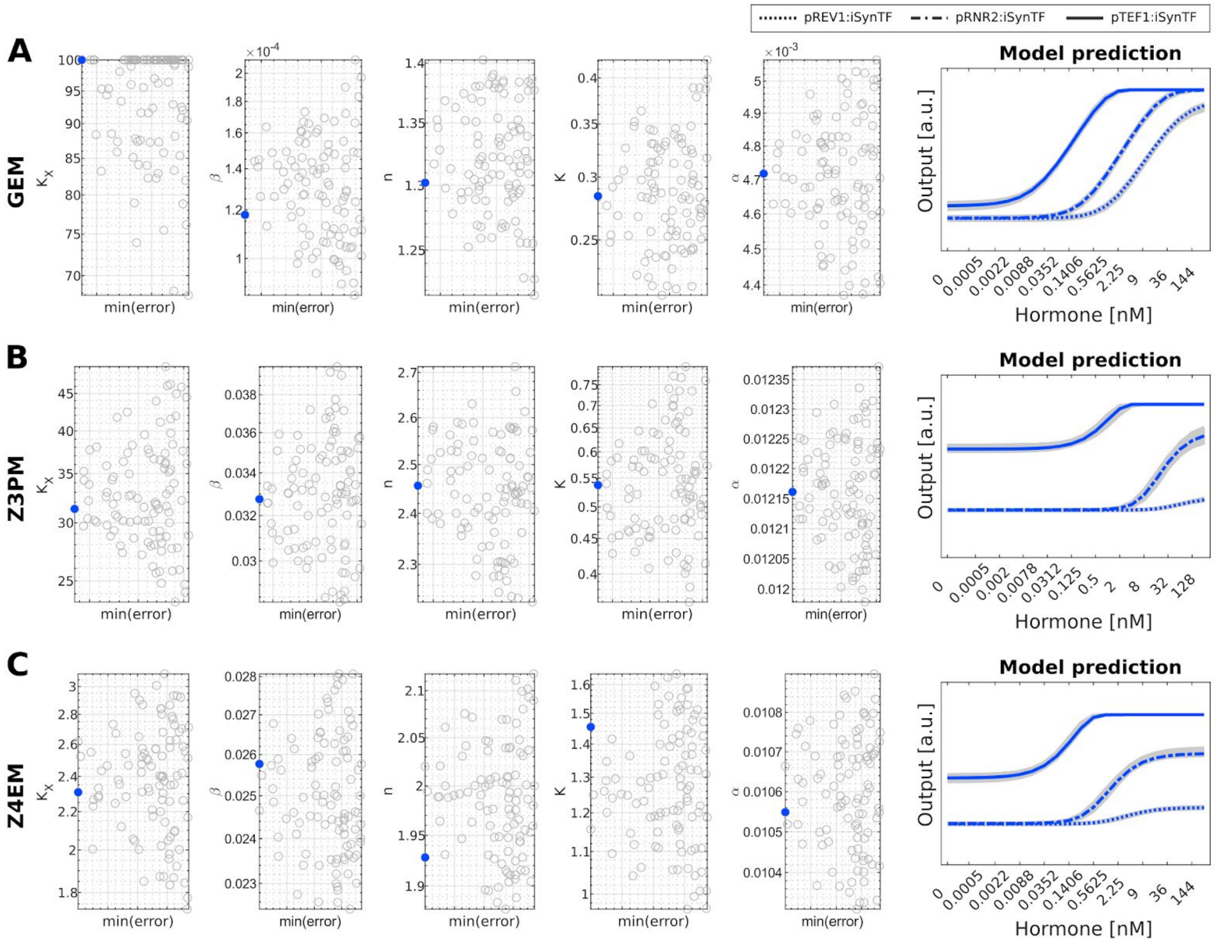


**Supplementary Figure 2: Results of parameter fitting of the mechanistic model to iSynTF dose-response data.** The top 100 (of 1000 tried) parameter sets in terms of their fitting error (see Methods) are shown for all fitted parameters for **(A)** GEM, **(B)** Z3PM, and **(C)** Z4EM**.** The mechanistic model predictions for each parameter set are shown at the right. For each plot, the best parameter set (i.e. lowest error) is highlighted in blue.

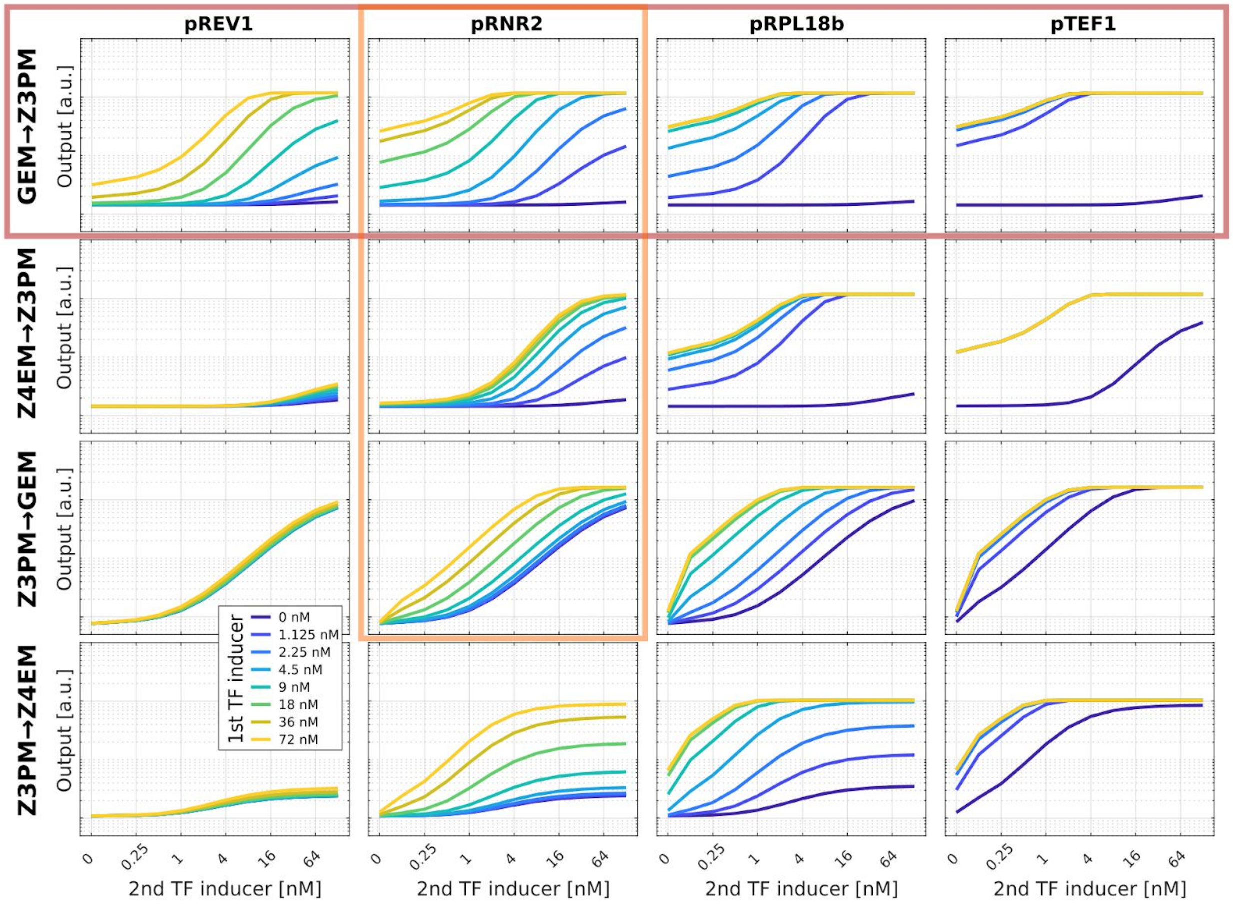


**Supplementary Figure 3: Using refined models to explore circuit designs.** For the proposed circuit in Fig. 3A, mechanistic model predictions for 2nd TF inducer dose responses (x-axis) for four possible arrangements varying the chosen iSynTFs (see row titles), at four different expression levels of the 1st TF expression level (see column titles), and 1st TF inducer concentrations (see legend). We experimentally verified six of these predictions, highlighted by the red (Fig.3B) and orange (Fig.3C) boxes here, and found substantial agreement.

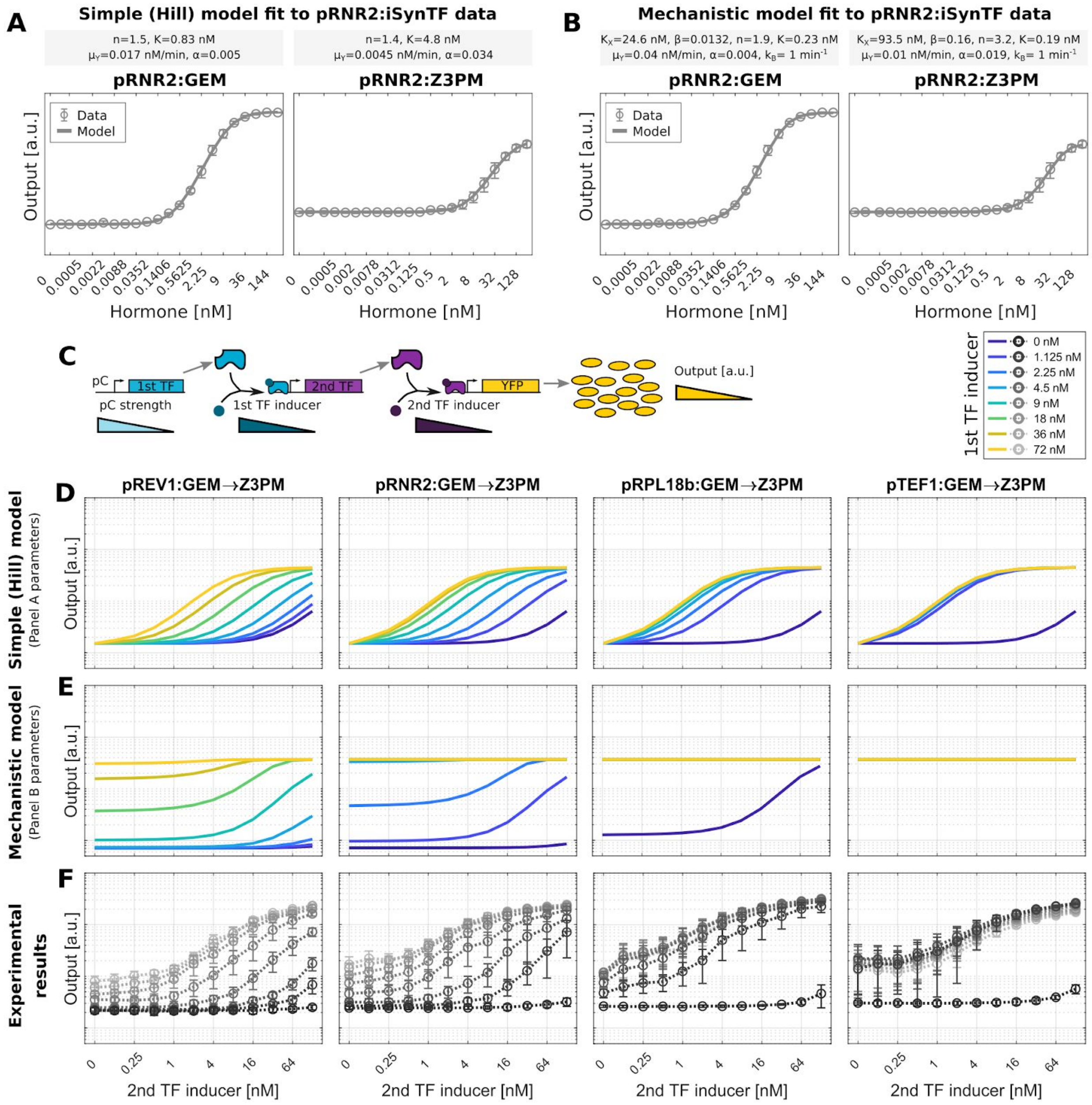


**Supplementary Figure 4: Models fit to a single hormone dose response fail to predict composability results.** Using only the inducer dose response data for GEM and Z3PM expressed from pRNR2, both the **(A)** *simple (Hill) model* (see Fig. 1A and Methods) and **(B)** *mechanistic model* (see Fig. 2A and Methods) can be fit equally well. The relevant parameter values are above each subpanel. **(C)** Diagram of the proposed circuit: a constitutively expressed iSynTF regulates the expression of a second iSynTF, which in turn regulates the expression of a reporter protein (output). Predicted response using the **(D)** *simple (Hill) model* and **(E)** *mechanistic model* fit to the pRNR2 data (above), compared to the **(F)** observed (experimental) response to changes in the 2nd TF inducer concentration (x-axis, Pg) as the constitutive promoter strength (see plot titles) and the 1st TF inducer concentration (legend) vary.

**Supplementary Table 1 - Integration plasmids used in this study (all KanR)**

| Name | Content | Yeast Auxotrophic Marker |
|---|---|---|
| pAN144 | pTEF1-GEM-tADH1 | ura3 |
| pAN145 | pRPL18b-GEM-tADH1 | ura3 |
| pAN146 | pRNR2-GEM-tADH1 | ura3 |
| pAN147 | pREV1-GEM-tADH1 | ura3 |
| pAN160 | pGal1-Venus-tPGK1 | leu2 |
| pAN248 | pTEF1-Z4EM-tADH1 | ura3 |
| pAN249 | pRPL18b-Z4EM-tADH1 | ura3 |
| pAN250 | pRNR2-Z4EM-tADH1 | ura3 |
| pAN251 | pREV1-Z4EM-tADH1 | ura3 |
| pAN360 | pTEF1-Venus-tPGK1 | leu2 |
| pAN361 | pRPL18b-Venus-tPGK1 | leu2 |
| pAN362 | pRNR2-Venus-tPGK1 | leu2 |
| pAN363 | pREV1-Venus-tPGK1 | leu2 |
| pGD240 | pZ4 (-Gal4 site)-Venus-tPGK1 | leu2 |
| pGD470 | pTEF1-GEM-tADH1 -- pGal1-Z3PM(fixed)-tPGK1 | ura3 |
| pGD471 | pRPL18b-GEM-tADH1 -- pGal1-Z3PM(fixed)-tPGK1 | ura3 |
| pGD472 | pRNR2-GEM-tADH1 -- pGal1-Z3PM(fixed)-tPGK1 | ura3 |
| pGD473 | pREV1-GEM-tADH1 -- pGal1-Z3PM(fixed)-tPGK1 | ura3 |
| pGD484 | pZ3-Venus-tSSA1 | leu2 |
| pGD485 | pZ3-Venus-tSSA1 -- pGal1-mScarlet-linker-tENO1 | leu2 |
| pGD488 | pTEF1-Z3PM(fixed)-tPGK1 | ura3 |
| pGD489 | pRPL18b-Z3PM(fixed)-tPGK1 | ura3 |
| pGD490 | pRNR2-Z3PM(fixed)-tPGK1 | ura3 |
| pGD491 | pREV1-Z3PM(fixed)-tPGK1 | ura3 |
| pGD532 | pGal1-Venus-tPGK1 -- pZ3-mScarlet-Linker-tENO1 | leu2 |
| pGD533 | pRNR2-Z3PM(fixed)-tPGK1 -- pZ3-GEM-tADH1 | ura3 |
| pGD898 | pRNR2-Z4EM-tADH1 -- pZ4 (-Gal4 site)-Z3PM(fixed)-tPGK1 | ura3 |

**Supplementary Table 2 - Oligos used in this study**

| Name | Sequence | Description |
|---|---|---|
| oAN019 | ACGCGGCCTTTTTACGGTTC | Dueber GG sequencing F |
| oAN074 | CTCCTGTTGATAGATCCAGT | Dueber GG sequencing R |
| g01 | GCATCGTCTCATCGGTCTCATATGAAGCTACTGTCTTCTATCGAACAAGCATGCGATATTTGCCGACTTAAAAAGCTCAAGTGCTCCAAAGAAAAACCGAAGTGCGCCAAGTGTCTGAAGAACAACTGGGAGTGTCGCTACTCTCCCAAAACCAAAAGaTCTCCGCTGACTAGGGCACATCTGACAGAAGTGGAATCAAGGCTAGAAAGACTGGAACAGCTATTTCTACTGATTTTTCCTCGAGAAGACCTTGACATGATTTTGAAAATGGATTCTTTACAGGATATAAAAGCATTGTTAACAGGATTATTTGTACAAGATAATGTGAATAAAGATGCCGTCACAGATAGATTGGCTTCAGTGGAGACTGATATGCCTCTAACATTGAGACAGCATAGAATAAGTGCGACATCATCATCGGAAGAGAGTAGTAACAAAGGTCAAAGACAGTTGACTGTATCGATTGACTCGGCAGGATCCGGTGACGGTGCTGGTTTAATTAACtctgctggagacatgagagctgccaacctttggccaagcccgctcatgatcaaacgctctaagaagaacagcctggccttgtccctgacggccgaccagatggtcagtgccttgttggatgctgagcccccatactctattccgagtatgatcctaccagacccttcagtgaagcttcgatgatgggcttactgaccaacctggcagacagggagctggttcacatgatcaactgggcgaagagggtgccaggctttgtggatttgaccctccatgatcaggtccaccttctagaatgtgcctggctaTGAGACGGCAT | GG3_GEM_gBlock_1 |

| | | |
|---|---|---|
| g02 | GCATCGTCTCAgctagagatcctgatgattggActcg<br>tctggcgctccatggagcacccagggaagctactgtt<br>tgctcctaacttgctcttggacaggaaccagggaaaa<br>tgtgtagagggcatggtggagatcttcgacatgctgc<br>tggctacatcatctcggttccgcatgatgaatctgca<br>gggagaggagtttgtgtgcctcaaatctattatttg<br>cttaattctggagtgtacacatttctgtccagcaccc<br>tgaagtctctggaagagaaggaccatatccaccgagt<br>cctggacaagatcacagacactttgatccacctgatg<br>gccaaggcaggcctgaccctgcaGCAGCAGCACCAGC<br>GGCTGGCCCAGCTCCTCCTCATCCTCTCCCACATCAG<br>GCACATGAGTAACAAAGGCATGGAGCATCTGTACAGC<br>ATGAAGTGCAAGAACGTGGTGCCCCTCTATGACCTGC<br>TGCTGGAGATGCTGGACGCCCACCGCCTACATGCGCC<br>CACTAGCCGTGGAGGGGCATCCGTGGAGGAaACGGAC<br>CAAAGCCACTTGGCCACTGCGGGCTCTACTTCATCGC<br>ATTCCTTGCAAAAGTATTACATCACGGGGGAGGCAGA<br>GGGTTTCCCTGCCACAGTCGCGGCtGCAGGTGACGGT<br>GCTGGTTTAATTAACATGACGGTCGACCATGATTTCA<br>ATAGCGAAGATATTTTATTCCCCATAGAAAGCATGAG<br>TAGTATACAATACGTGGAGAATAATAACCCAAATAAT<br>ATTAACAACGATGTTATCCCGTATTCTCTAGATATCA<br>ATGAGACGGCAT | GG3_GEM_gBlock_2 |
| g03 | GCATCGTCTCATCAAAAACACTGTCTTAGATAGTGCG<br>GATCTCAATGACATTCAAAATCAAGAAACTTCACTGA<br>ATTTGGGGCTTCCTCCACTATCTTTCGACTCTCCACT<br>GCCCGTAACGGAAACGATACCATCCACTACCGATAAC<br>AGCTTGCATTTGAAAGCTGATAGCAACAAAAATCGCG<br>ATGCAAGAACTATTGAAAATGATAGTGAAATTAAGAG<br>TACTAATAATGCTAGTGGCTCTGGGGCAAATCAATAC<br>ACAACTCTTACTTCACCTTATCCTATGAACGACATTT<br>TGTACAACATGAACAATCCGTTACAATCACCGTCACC<br>TTCATCGGTACCTCAAAATCCGACTATAAATCCTCCC<br>ATAAATACAGCAAGTAACGAAACTAATTTATCGCCTC<br>AAACTTCAAATGGTAATGAAACTCTTATATCTCCTCG<br>AGCCCAACAACATACGTCCATTAAAGATAATCGTCTG<br>TCCTTACCTAATGGTGCTAATTCGAATCTTTTCATTG<br>ACACTAACCCAAACAATTTGAACGAAAAACTAAGAAA<br>TCAATTGAACTCAGATACAAATTCATATTCTAACTCC<br>ATTTCTAATTCAAACTCCAATTCTACGGGTAATTTAA<br>ATTCCAGTTATTTTAATTCACTGAACATAGACTCCAT<br>GCTAGATGATTACGTTTCTAGTGATCTCTTATTGAAT<br>GATGATGATGATGACACTAATTTATCACGCCGAAGAT<br>TTAGCGACGTTATAACAAACCAATTTCCGTCAATGAC<br>AAATTCGAGGAATGAGCTCGGATCCTGAGACCTGAGA<br>CGGCAT | GG3_GEM_gBlock_3 |

| | | |
|---|---|---|
| oAN030 | GCATCGTCTCATCGGTCTCATATGGGTACCCGCCCATATG | GG3_Z3PM/Z4EM_PCR_1_F |
| oAN145 | ATGCCGTCTCACTGCAGCCGCCTTTTTATGAAAGAG | GG3_Z3PM_PCR_1_R |
| oAN146 | GCATCGTCTCAGCAGGTGACGGTGCTGGTTTAAT | GG3_Z3PM_PCR_2_F |
| oAN147 | ATGCCGTCTCAGGTCTCAGGATCCGAGCTCTTGGTTTGTTATAACG | GG3_Z3PM(fixed)_PCR_2_R |
| oAN027 | ATGCCGTCTCACAATCATCAGGATCTCTAGCCAG | GG3_Z4EM_PCR_1_R |
| oAN028 | GCATCGTCTCAATTGGACTCGTCTGGCGCTCC | GG3_Z4EM_PCR_2_F |
| oAN031 | ATGCCGTCTCAGGTCTCAGGATCCGAGCTCATTCCTCGAATTTG | GG3_Z4EM_PCR_2_R |
| oAN022 | GCATCGTCTCATCGGTCTCAAACGTTATATTGAATTTTCAAAAATTCTTACTTTTTTTTT | GG2_pZ3/4_PCR_1_F |
| oAN025 | ATGCCGTCTCAGGTCTCACATAGATCTTATAGTTTTTTCTCCTTGACGTTAAAG | GG2_pZ3_PCR_1_R |
| oGD122 | ATGCCGTCTCAGCAattTCTAGACTCCTCCGCC | GG2_pZ4_(-Gal4_site)_PCR_1_R |
| oGD123 | GCATCGTCTCAtTGCGTCCTCGTCTTCACCG | GG2_pZ4_(-Gal4_site)_PCR_2_F |
| oGD124 | ATGCCGTCTCAGGTCTCACATAGATCTGATCTTATAGTTTTTTCTCCTTGAC | GG2_pZ4_(-Gal4_site)_PCR_2_R |

**Supplementary Table 3 - Yeast strains used in this study**

| Strain | Genotype | Background Strain | Plasmid(s) |
|---|---|---|---|
| yWCD230 | BY4741, HIS3 repaired (MATa, his3Δ1, leu2Δ0, met15Δ0, ura3Δ0) | | |
| yAHN061 | pGal1-Venus-tPGK1::leu2; BY4741, HIS3 repaired | yWCD230 | pAN160 |
| yAHN066 | pTEF1-GEM-tADH1::ura3; pGal1-Venus-tPGK1::leu2; BY4741, HIS3 repaired | yAHN061 | |
| yAHN067 | pRPL18B-GEM-tADH1::ura3; pGal1-Venus-tPGK1::leu2; BY4741, HIS3 repaired | yAHN061 | |
| yAHN068 | pRNR2-GEM-tADH1::ura3; pGal1-Venus-tPGK1::leu2; BY4741, HIS3 repaired | yAHN061 | |
| yAHN069 | pREV1-GEM-tADH1::ura3; pGal1-Venus-tPGK1::leu2; BY4741, HIS3 repaired | yAHN061 | |
| yGD074* | pTEF1-mTagBFP2-tENO2::HO, HIS3; pTEF1-mScarlet-tADH1::ura3; pTEF1-Venus-tPGK1::leu2; BY4741, HIS3 repaired | yWCD230 | pGD151; pAN548; pAN360 |
| yGD075* | pRPL18b-mTagBFP2-tENO2::HO, HIS3; pRPL18b-mScarlet-tADH1::ura3; pRPL18b-Venus-tPGK1::leu2; BY4741, HIS3 repaired | yWCD230 | pGD152; pAN434; pAN361 |
| yGD076* | pRNR2-mTagBFP2-tENO2::HO, HIS3; pRNR2-mScarlet-tADH1::ura3; pRNR2-Venus-tPGK1::leu2; BY4741, HIS3 repaired | yWCD230 | pGD153; pAN549; pAN362 |
| yGD077* | pREV1-mTagBFP2-tENO2::HO, HIS3; pREV1-mScarlet-tADH1::ura3; pREV1-Venus-tPGK1::leu2; BY4741, HIS3 repaired | yWCD230 | pGD154; pAN550; pAN363 |
| yGD175 | pZ4 (-Gal4 site)-Venus-tPGK1::leu2; BY4741, HIS3 repaired | yWCD230 | pGD240 |
| yGD177 | pTEF1-Z4EM-tADH1::ura3; pZ4 (-Gal4 site)-Venus-tPGK1::leu2; BY4741, HIS3 repaired | yGD175 | pAN248 |
| yGD178 | pRPL18B-Z4EM-tADH1::ura3; pZ4 (-Gal4 site)-Venus-tPGK1::leu2; BY4741, HIS3 repaired | yGD175 | pAN249 |
| yGD179 | pRNR2-Z4EM-tADH1::ura3; pZ4 (-Gal4 site)-Venus-tPGK1::leu2; BY4741, HIS3 repaired | yGD175 | pAN250 |
| yGD180 | pREV1-Z4EM-tADH1::ura3; pZ4 (-Gal4 site)-Venus-tPGK1::leu2; BY4741, HIS3 repaired | yGD175 | pAN251 |
| yGD307 | pZ3-Venus-tSSA1::leu2; BY4741, HIS3 repaired | yWCD230 | pGD484 |
| yGD308 | pZ3-Venus-tSSA1 -- pGal1-mScarlet-Linker-tENO1::leu2; BY4741, HIS3 repaired | yWCD230 | pGD485 |

| | | | |
|---|---|---|---|
| yGD310 | pTEF1-Z3PM(fixed)-tPGK1::ura3; pZ3-Venus-tSSA1::leu2; BY4741, HIS3 repaired | yGD307 | pGD488 |
| yGD311 | pRPL18B-Z3PM(fixed)-tPGK1::ura3; pZ3-Venus-tSSA1::leu2; BY4741, HIS3 repaired | yGD307 | pGD489 |
| yGD312 | pRNR2-Z3PM(fixed)-tPGK1::ura3; pZ3-Venus-tSSA1::leu2; BY4741, HIS3 repaired | yGD307 | pGD490 |
| yGD313 | pREV1-Z3PM(fixed)-tPGK1::ura3; pZ3-Venus-tSSA1::leu2; BY4741, HIS3 repaired | yGD307 | pGD491 |
| yGD325 | pTEF1-GEM-tADH1 -- pGal1-Z3PM(fixed)-tPGK1::ura3; pZ3-Venus-tSSA1 -- pGal1-mScarlet-Linker-tENO1::leu2; BY4741, HIS3 repaired | yGD308 | pGD470 |
| yGD326 | pRPL18B-GEM-tADH1 -- pGal1-Z3PM(fixed)-tPGK1::ura3; pZ3-Venus-tSSA1 -- pGal1-mScarlet-Linker-tENO1::leu2; BY4741, HIS3 repaired | yGD308 | pGD471 |
| yGD327 | pRNR2-GEM-tADH1 -- pGal1-Z3PM(fixed)-tPGK1::ura3; pZ3-Venus-tSSA1 -- pGal1-mScarlet-Linker-tENO1::leu2; BY4741, HIS3 repaired | yGD308 | pGD472 |
| yGD328 | pREV1-GEM-tADH1 -- pGal1-Z3PM(fixed)-tPGK1::ura3; pZ3-Venus-tSSA1 -- pGal1-mScarlet-Linker-tENO1::leu2; BY4741, HIS3 repaired | yGD308 | pGD473 |
| yGD354 | pGal1-Venus-tENO2 -- pZ3-mScarlet-Linker-tENO1::leu2; BY4741, HIS3 repaired | yWCD230 | pGD532 |
| yGD356 | pRNR2-Z3PM(fixed)-tPGK1 -- pZ3-GEM-tADH1::ura3; pGal1-Venus-tENO2 -- pZ3-mScarlet-Linker-tENO1::leu2; BY4741, HIS3 repaired | yGD354 | pGD533 |
| yGD588 | pRNR2-Z4EM-tADH1 -- pZ4 (-Gal4 site)-Z3PM(fixed)-tPGK1::ura3; pZ3-Venus-tSSA1::leu2; BY4741, HIS3 repaired | yGD307 | pGD898 |

* Plasmids represent BFP, RFP, and YFP cassettes. Each was integrated separately in the reverse order. Only the YFP plasmid is listed in Supplementary Table 1 as it was the only measurement taken.

**pZ4 (-Gal4 site) sequence**

TTATATTGAATTTTCAAAAATTCTTACTTTTTTTTTTGGATGGACGCAAAGAAGTTTAATAATCA
TATTACATGGCATTACCA***CCATATACATAT***CCATATACATATCCATATCTAATCTTACTTATAT
GTTGTGGAAATGTAAAGAGCCCCATTATCTTAGCCTAAAAAAACCTTCTCTTTGGAACTTTCAG
TAATACGCTTAACTGCTCATTGCTATATTGAAGTGTGGCC**GCGGCGGAGGAG**T**GCGGCGGAGGA**
**G**GA**GCGGCGGAGGAG**T**GCGGCGGAGGAG**GA**GCGGCGGAGGAG**T**GCGGCGGAGGAG**TCTAGA<u>**AAT**</u>
TGCGTCCTCGTCTTCACCGGTCGCGTTCCTGAAACGCAGATGTGCCTAATGCCGCACTGCTCCG
AACAATAAAGATTCTACAATACTAGCTTTTATGGTTATGAAGAGGAAAAATTGGCAGTAACCTG
GCCCCACAAACCTTCAAATTAACGAATCAAATTAACAACCATAGGATGATAATGCGATTAGTTT
TTTAGCCTTATTTCTGGGGTAATTAATCAGCGAAGCGATGATTTTTGATCTATTAACAGATATA
TAAATGGAAAAGCTGCATAACCACTTTAACTAATACTTTCAACATTTTCAGTTTGTATTACTTC
TTATTCAAATGTCATAAAAGTATCAACAAAAAATTGTTAATATACCTCTATACTTTAACGTCAA
GGAGAAAAAACTATA***AGATC***

**Bold:** Z4 operator sequence
<u>**Bold and underline**</u>: CCG mutated to AAT to break Gal4 consensus site
***Bold and Italicized***: insertions from cloning. Upstream insertion sequence seems to be a duplication from amplification. Terminal insertion sequence was a copy error due to implementing YTK part standards.
